# Visual control of landing maneuvers in houseflies on vertical and inverted surfaces

**DOI:** 10.1101/448472

**Authors:** Sujay Balebail, Satish K Raja, Sanjay P. Sane

## Abstract

Landing maneuvers in flies are complex behaviors that may be conceptually decomposed into a sequence of modular behaviors such as body deceleration, extension of legs, and body rotations which are coordinated to ensure controlled touchdown. The composite nature of these behaviors means that there is variability in the kinematics of landing maneuvers, making it difficult to identify the general rules that govern this behavior. Many previous studies have relied on tethered preparations to study landing behaviors, but tethering constrains some behavioral modules to operate in an open feedback control loop while others remain in closed-loop, thereby inducing experimental artefacts. On the other hand, freely flying insects are hard to precisely control, which may also increase behavioral variability. One approach towards understanding the general rules underlying landing behavior is to determine the common elements of landing kinematics on surfaces that are oriented in different ways. We conducted a series of experiments in which the houseflies, *Musca Domestica*, were lured to specific visual targets on either vertical or inverted horizontal substrates. These conditions elicited landing behaviors in the flies that could be captured accurately using multiple high-speed video cameras. We filmed the houseflies landing on surfaces oriented along two directions: vertical (*vertical landings*), and upside down (*inverted landings*). Our experiments reveal that flies that are able to land feet-first in a controlled manner must satisfy specific criteria, failing which their landing performance is compromised causing their heads to bump into the surface during landing. Flies landing smoothly on both surfaces initiate deceleration at approximately fixed distances from the substrate and in direct proportion to the component of flight velocity normal to the landing surface. The ratio of perpendicular distance to the substrate and velocity at the onset of deceleration was conserved, despite the large differences in the mechanics of the vertical vs. inverted landings. Flies extend their legs independently of distance from the landing surface or their approach velocity normal to the surface, regardless of the orientation of the landing substrate. Together, these results show that the visual initiation of deceleration is robust to orientation of the landing surface, whereas the initiation of leg-extension may be context-dependent and variable which allows flies to land on substrates of various orientations in a versatile manner. These findings may also be of interest to roboticists that are interested in developing flapping robots that can land on surfaces of different orientations.

## INTRODUCTION

Safe landing on a substrate is a key aspect of insect flight behavior. In their natural world, the surfaces on which insects land are oriented in diverse ways, and hence the underlying behavioral principles that guide their landing behavior must enable such versatility. From the controls’ perspective, smooth landing requires insects to rapidly sense and precisely react to an approaching substrate in a manner that is robust to diverse orientations the landing surface. It has been previously suggested that landing behavior can be subdivided into many distinct, independently-activated behaviors, and may therefore be considered as ‘modular’ (van Breugel and Dickinson, 2012). While landing, insects typically reduce their approach velocities (Baird et al., 2013; Lee et al., 1991; Lee et al., 1993; Srinivasan et al., 2000; van Breugel and Dickinson, 2012; Wagner, 1982), extend their legs (Goodman, 1960; Evangelista et al., 2010; Hyzer, 1962; Lee et al., 1993; Reber et al., 2016a; Reber et al., 2016b; van Breugel and Dickinson, 2012), and align their body parallel the landing surface (Hyzer, 1962; Zhao et al., 2017). Moreover, insects land on objects of different textures, flexibility, and orientations including inverted surfaces (Evangelista et al., 2010; Hyzer, 1962; Reber et al., 2016a), suggesting a great degree of adaptability of their landing behavior.

What basic strategies underlie the versatile landing ability of insects? To address this question, we must consider the following key points. First, any strategy to initiate deceleration must ensure that the animal has sufficient time to achieve low contact velocities, thereby avoiding injuries upon impact. The rules used to determine the onset of deceleration have been studied in freely-flying houseflies *Musca domestica* (Wagner, 1982) and fruit flies *Drosophila melanogaster* (van Breugel and Dickinson, 2012). An important parameter in these studies is the ratio of the distance of the flying insect from the landing object and the velocity component in the direction of the object, which is conventionally termed *tau* (e.g. Lee, 1980 and associated discussion by Kalmus; also Baird et al, 2013). The value of *tau* at any time instant represents the time to collision with the landing surface, as the animal flies towards the landing object. Wagner (1982) showed that houseflies approaching a spherical landing object initiated deceleration when the value of *tau* fell below a threshold value. Thus, flies approaching an object at higher velocities initiated deceleration proportionately further away from the object i.e. at a constant value of *tau.*

To a landing fly, the main sensory cues that are available are the rates of optic flow on their retina, which indicate how fast the object is approaching the fly. Accounting for this, Wagner et al (1982) proposed the Relative Retinal Expansion Velocity (RREV) model, which suggests that flies initiate deceleration at a critical value of the ratio of retinal expansion velocity to the retinal size of an object. Another model called the Retinal Size-Dependent Expansion Threshold (RSDET) Model was proposed to explain the data on landing maneuvers in *Drosophila melanogaster* (van Breugel and Dickinson, 2012). Specifically, their instantaneous approach speed was proportional to the logarithm of the angular size subtended by the post on the retina. The RSDET Model (van Breugel and Dickinson, 2012) specifically addressed the onset of deceleration in *Drosophila* as they approached a cylindrical post, and proposed that deceleration is initiated at a threshold value of the retinal size dependent expansion of the object on the retina. How fast the fly can cross this threshold depends on the its speed of approach, but not on the physical dimensions of the object. Thus, a small object that expands slowly is as likely to trigger onset of deceleration as a large object that expands rapidly. Similarly, a fly that is further away from flying faster would initiate deceleration as would a fly that is flying slowly but is closer to the substrate. In most practical matters, the RSDET model is similar to the RREV or tau-estimation models.

While landing, the rate of deceleration needs to be controlled to achieve smooth touchdown. Birds such as hummingbirds (Lee et al., 1991), and pigeons (Lee et al., 1993) control deceleration by maintaining the rate of change of *tau* with time at a constant value between 0.5 and 1. Honeybees, on the other hand, keep *tau* at a fixed value after initiating deceleration, ensuring that the component of flight velocity normal to the landing surface reduces linearly with displacement from the surface (Baird et al., 2013; Srinivasan et al., 2000). Freely flying insects extend their legs before contacting the surface (Evangelista et al., 2010; Hyzer, 1962; Lee et al., 1993; Reber et al., 2016a; Reber et al., 2016b; van Breugel and Dickinson, 2012). The rules governing the initiation of the leg-extension response in free flight has been the subject of many previous studies (Goodman, 1960; Evangelista et al., 2010; Lee et al., 1993; Reber et al., 2016a; van Breugel and Dickinson, 2012; Baird et al, 2013). For instance, pigeons approaching a perch to land, begin extending their legs at a fixed value of *tau* (Lee et al., 1993). When honeybees (Evangelista et al., 2010) and bumblebees (Reber et al., 2016a) approached plane surfaces, they were observed to hover and extend their legs at a constant distance from the landing surface, irrespective of the inclination of the surface. When *Drosophila melanogaster* approached a cylindrical post, the onset of leg-extension appeared to be independent of approach velocity, depending instead on a threshold distance from the post or threshold angle subtended by the post on the retina (van Breugel and Dickinson, 2012).

These modules can be independently activated; for example, presentation of front-to-back optic flow stimuli to tethered insects elicits a leg-extension response (Borst, 1986; Borst, 1989; Borst and Bahde, 1986, 1987, 1988a, b, and 1990; Coggshall, 1972; De Talens and Ferreti, 1970; Eckert, 1980; Goodman, 1960; Tammero and Dickinson, 2002) even though there is no physical deceleration or change in body pitch. This behavior is thought to be analogous to a freely-flying insect extending its legs before touchdown to prevent a crash landing. In tethered houseflies, the time course of leg-extension remains fairly constant regardless of the nature of the releasing stimulus. However, the latency of the leg-extension response depends on the optic flow stimulus (Borst, 1986), and is a function of the size, velocity, and contrast of looming stimuli (Borst, 1990; Borst and Bahde, 1988a; Goodman, 1960). Besides extending their legs, tethered flies also reduce their thrust in response to a looming stimulus, and the onsets of reduction in thrust is correlated with leg-extension (Borst and Bahde, 1988a).

Despite the extensive research on landing responses, several questions have remained largely unanswered that relate to the mutual coordination between leg extension and deceleration of the body. Do flies follow certain rules for the initiation of these two modules of landing? Are these rules dependent on the orientation of the landing surface? Are these modules initiated using the same rules, or are they initiated independently? To address these questions, it is necessary to determine the generalities of landing responses, irrespective of orientation of landing. Here, we filmed at high frame rates (3000 or 4000 fps), the landing behavior of houseflies *(Musca domestica)* on plane surfaces oriented along two directions, vertical (vertical landings) and upside down (inverted landings).

Flies approaching the landing surface at higher velocities must slow down at an appropriate distance from the surface and also extend their legs to avoid injuries upon impact. We hypothesized that this imposes on them the need for coordination between body deceleration and leg extension responses. Moreover, such coordination is required regardless of the contexts in which these behaviors occur. Previous free-flight studies indicated that flying insects begin leg-extension at a fixed displacement from the landing surface in which case the inter-trial variability in displacement at the beginning of leg-extension is expected to be low. Tethered flight studies in houseflies indicate that the onsets of deceleration and leg-extension are correlated (Borst and Bahde, 1988a), implying that the similar visual cues initiate both responses, however with different latencies. If so, we expect a fixed time difference between the onsets of deceleration and leg-extension.

## MATERIALS AND METHODS

### Animals

Adult houseflies *(Musca domestica)* were captured from the wild and stored in a container with ad libitum access to sucrose and water.

### Experimental setup and protocol

#### Vertical landings

The flight chamber for filming vertical landings comprised of a transparent plexiglass box (28 cm × 28 cm × 28 cm). Three 4.5 cm × 4.5 cm pieces of chart paper were attached to form an equilateral prism-shaped object and its edges were lined with black strips. This object was placed approximately in the center of the chamber, and served as the landing substrate for the fly. The chamber was lit by a studio light (Simpex Compact 300, Simpex Industries, Delhi, India) to ∼3000 lux (measured using a Center 337 light meter, Center Technology Corporation, Taipei, Taiwan). Flies were introduced into the filming chamber from the top. Landings on the object were recorded at 3000 fps by two synced high speed cameras (Phantom v7.3, Vision Research, Wayne, NJ, USA; Fig 1 A, Ai). The field of view of both high-speed cameras were calibrated using a standard object. The flies generally performed a saccade towards the object before landing, as has also been reported in the case of *Drosophila melanogaster* (van Breugel and Dickinson, 2012). The frame where the saccade appeared to end was selected as the start point of each video. The frame of first contact with the landing surface was chosen as the end point of the video.

**Fig 1.**
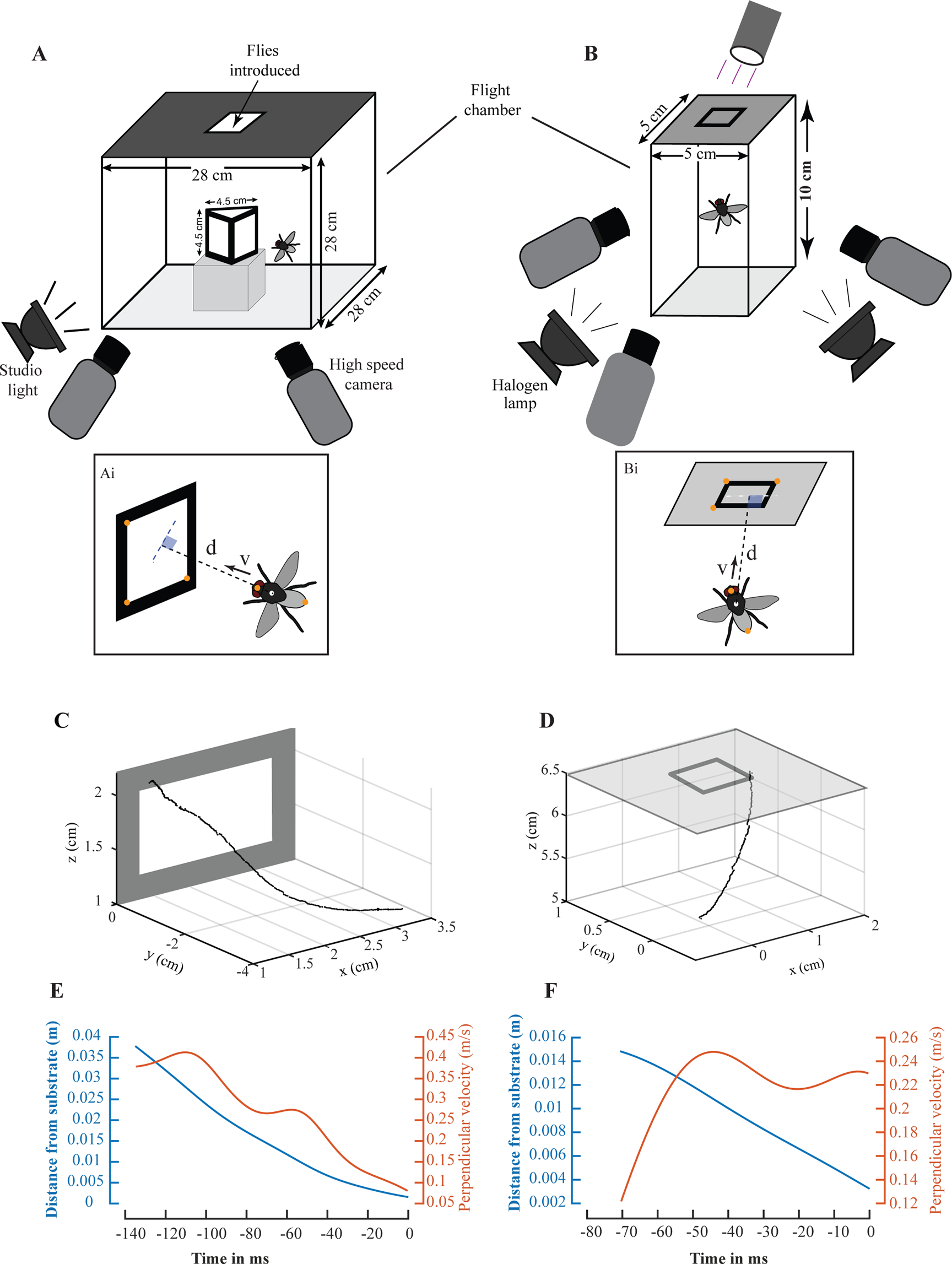
Experimental setups to record vertical and inverted landings, and measurement of the associated flight variables. (A) The experimental setup for filming vertical landings which were elicited on a prism-shaped object and recorded by two synchronized high-speed cameras at 3000 fps. (B) Experimental setup for filming inverted landings which occurred on a translucent ceiling and recorded by three synchronized high-speed cameras at 4000 fps. For both vertical landings (Ai) and inverted landings (Bi), we digitized the tips of the head and abdomen of the fly in each frame, in addition to three points on the landing surface. We computed the midpoint of the line joining the head and abdomen tips, and the distance of the midpoint from the landing surface (d). The component of flight velocity perpendicular to the plane of the landing surface (v) was computed according to Equation 1. (C-D) Sample raw trajectories of the midpoint of a fly performing a vertical (C) and inverted landing (D). (E-F) Below each trajectory, the distance from the substrate (blue trace) and perpendicular velocity (orange trace) are plotted as functions of time to collision to the landing surface. The flies contacted the landing surface at 0 ms.

#### Inverted landings

The flight chamber for filming inverted landings comprised of a glass box (5 cm × 5 cm × 10 cm) with a translucent filter paper ceiling (Fig 1B). A black square outline (side length= 1.5 cm, line thickness= 2 mm) was drawn approximately on the center of the ceiling, to provide an expansion stimulus as the fly approached the ceiling. A batch of 3-6 flies were starved for 10-12 hours, anesthetized *via* a 2.5 min cold shock (−20°C) and placed in the filming chamber. The chamber was illuminated by a UV torch placed above the ceiling (to attract flies), two 150 W halogen lamps, and two stereomicroscope lights (Nikon SMZ25; Nikon Corporation, Tokyo, Japan), to ∼30000 lux. The anesthetized flies were allowed to recover for 10-15 minutes. Landings on the ceiling were recorded by three synced high-speed cameras filming at 4000 fps (two phantom v7.3 and one phantom v611; Fig. 1B, Bi). No more than one landing was recorded per batch of flies, to avoid pseudo-replication. The field of view of the three cameras were calibrated using a standard object. In most trials, the fly took off from a lateral wall, rotated about its longitudinal axis (roll rotation) by almost 360°, and then ascended towards the ceiling. The frame in which the roll rotation ended was determined by a careful observation of the recording. It was chosen as the as the start point and the frame of first contact with the landing surface was selected as the end point of each video.

#### Digitization and computation of flight variables

Videos of landings were digitized using custom MATLAB software (Hedrick, 2008; Mathworks, Natick, MA, USA). We digitized the tips of the head and abdomen, and three points on the landing surface (Fig. 1Ai, Bi). The time series of the digitized points was filtered using a 4th order low-pass filter (Butterworth) with a cut-off frequency 30 Hz. This was done to eliminate the influence of body rotations, whose mean frequency was 50±23 (μ±σ) Hz (Fig. S1A). Before applying the filter, the ends of the time series data were extrapolated using quadratic functions to reduce edge effects (Walker, 1998). The coordinates of the midpoint of the line joining the head and abdomen tips was computed (henceforth termed “midpoint”) at each frame to determine the broad trajectories during landing (Fig 1 C, D). Two flight variables were computed from the digitized points: First, the perpendicular (shortest) distance of the midpoint from the landing surface (d) and second, the component of flight velocity perpendicular to the plane of the landing surface (v), for each frame in the following manner:

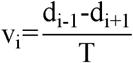

in which the subscript i stands for the frame number and T is the time interval between (i-1)th and (i+1) frames (2/3 ms for vertical landings and 1/2 ms for inverted landings (Fig 1 E, F)).

#### Identifying the onsets of deceleration and leg-extension

We wrote custom code in MATLAB to identify all the local maxima and minima in the plots of perpendicular velocity (v) *vs* time (Fig. 2A-B, E-F) and distance from substrate as a function of perpendicular velocity (Fig 2 C-D, G-H). Trials in which the final extremum before touchdown was a minimum were classified as having no deceleration before touchdown (Fig. 2B, F). In the remaining trials, the final maximum velocity before first contact with the landing surface was classified as the onset of deceleration (Fig. 2A, E).

**Fig 2.**
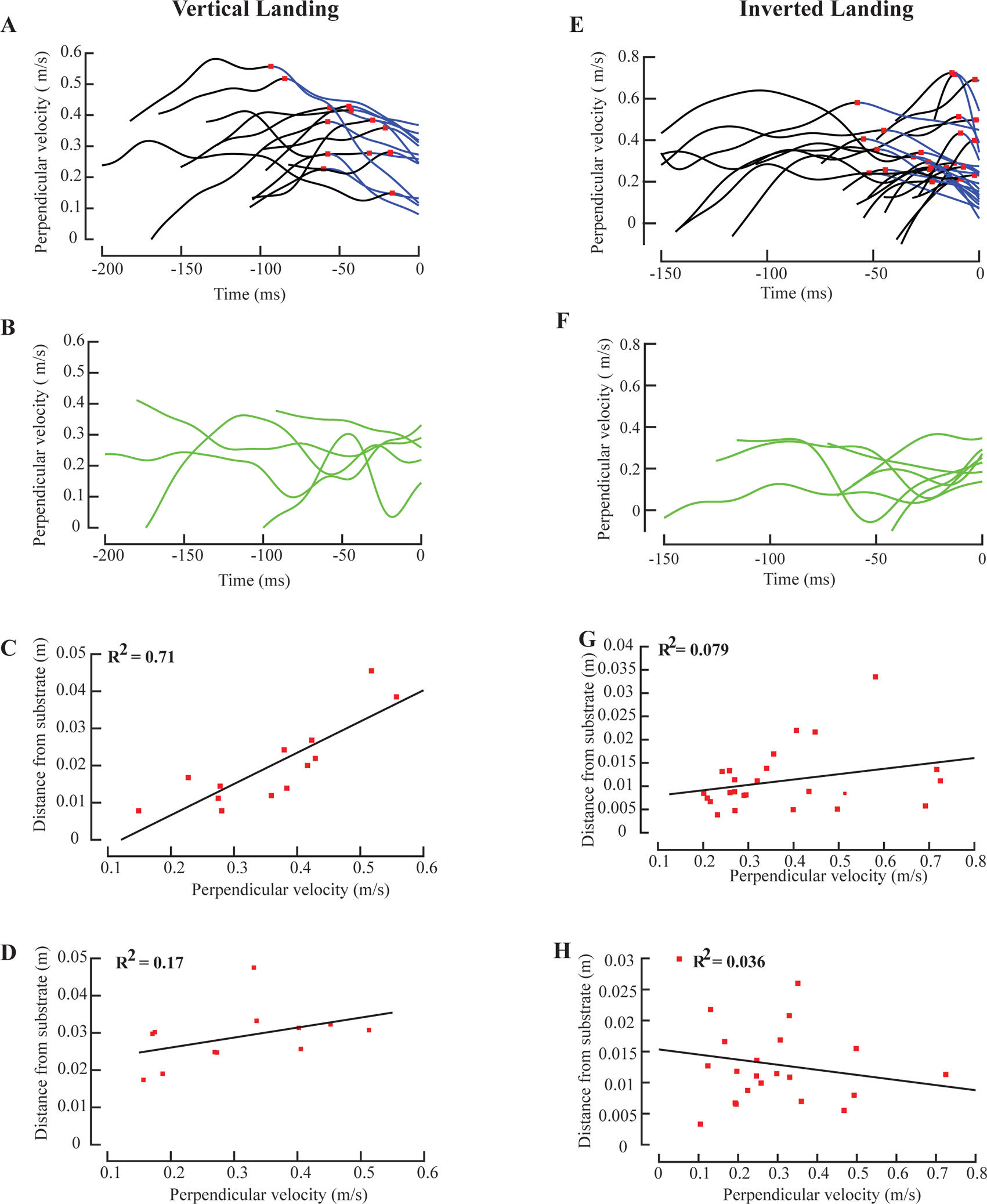
Initiation of deceleration and leg-extension during vertical and inverted landings. (A) Perpendicular velocity versus time to collision for all vertical landing trials in which the fly decelerated before touchdown on the vertical surface (n=13). The flies contacted the landing surface at 0 ms. The red squares mark the onset of deceleration as identified by our code (see Materials and methods). The blue sections of the traces represent the decelerating segments of the flight trajectory. (B) Perpendicular velocity versus time to collision for the vertical landing trials in which the fly did not decelerate before touchdown (n=5). (C) Distance from substrate versus perpendicular velocity at the time of onset of deceleration for the 13 vertical landing trials. The coefficient of determination (R^2^) of the best fit line is 0.71. (D) Distance from substrate versus perpendicular velocity at the onset of leg-extension for the 12 vertical landings whose frame of onset of leg-extension could be identified (see Materials and methods). R^2^ of the best fit line is 0.17. (E) Perpendicular velocity versus time for all inverted landing trials in which the fly decelerated before touchdown on the upside down surface (n=25). (F) Perpendicular velocity versus time for the inverted landing trials in which the fly did not decelerate before touchdown (n=7). (G) Distance from substrate versus perpendicular velocity at the onset of deceleration for 25 inverted landing trials. R^2^ of the best fit line is 0.079. (H) Distance from substrate versus perpendicular velocity at the onset of leg-extension for the 22 inverted landing trials in which the flies extended their legs while approaching the landing surface (and not during take-off, see Materials and methods). R^2^ of the best fit line is 0.036.

Unlike the onset of deceleration which required the above calculations, the onset of leg-extension could be visually determined from a close examination of the videos. The frame in which either one or both the front legs began to be raised dorsally was chosen as the frame of onset of leg-extension. In 6 out of the 18 vertical landing trials, the fly had extended its legs before arriving in the field of view of both cameras. Therefore, the frame of onset of leg-extension is unknown for these trials. In 10 out of 32 inverted landing trials, the fly extended its legs at the takeoff point and kept them extended. For these trials, leg extension could not be attributed to landing *per se,* and hence we did not include these trials in the analysis of the initiation of leg extension.

#### Testing hypotheses for the initiation of deceleration and leg-extension

To test whether flies initiate both components of the landing behavior at a distance that is proportional to perpendicular velocity (constant *tau* hypothesis), we plotted distance from the substrate (d) against perpendicular velocity (v) at the onsets of deceleration (Fig. 2 C, G; Fig. 3 C-D; Fig. 5 A-B) and leg-extension (Fig. 2 D, H; Fig. 4 A-B), and computed the coefficient of determination (R^2^) of the best fit line using in-built functions in MATLAB. The slope of this best-fit line is defined as *tau.* High R^2^ values would support the constant *tau* hypothesis. If flies initiate a module at a fixed distance from the landing platform, then the inter-trial variability in distance at the onset of the module is expected to be low. If flies utilize the same cues for releasing both deceleration and leg-extension but with different latencies, we should expect stereotypy in the time difference between the modules. We next plotted time to collision to the landing surface at the onset of leg-extension (time difference between the onset of leg-extension and first contact with the landing surface) as a function of time to collision to the landing surface at the onset of deceleration (time difference between the onset of deceleration and first contact with the landing surface; Fig. 5 E-F), and fit lines to the plots. High R^2^ values would support the hypothesis of both modules being initiated by the same stimuli.

**Fig 3.**
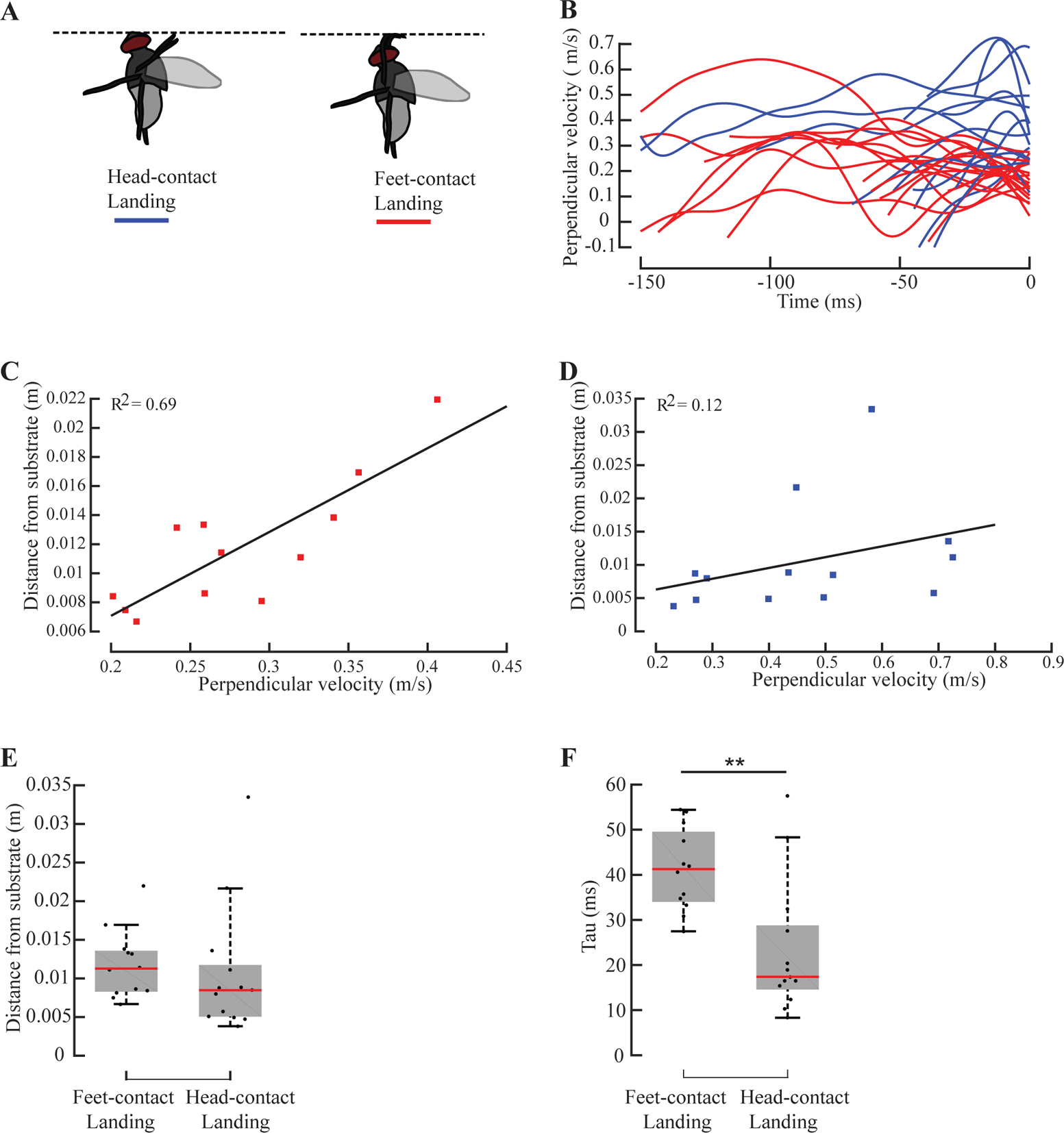
Initiation of deceleration for flies performing feet-contact and head-contact inverted landings. (A) A closer look at the videos of inverted landings revealed that the trials can be grouped into two categories. In 15 trials, the head made contact with the landing surface (“Head-contact landings”, blue) whereas in the remaining 17 trials, the head did not touch the landing surface during the course of the landing maneuver (“feet-contact landings”, red) (B) Perpendicular velocity versus time to collision for all trials (n=32) (C-D) Only 25 out of the 32 flies decelerated before landing (see Materials and methods), and analyzed further. Of these 25 inverted landing trials, 12 flies performed a feet-contact landing and 13 flies executed a head-contact landing. (C) Distance from substrate versus perpendicular velocity at the time of onset of deceleration for inverted feet-contact landing trials (n=12, R^2^ = 0.69). (D) Distance from substrate versus perpendicular velocity at the time of onset of deceleration for inverted head-contact landing trials (n=13, R^2^ = 0.12). (E-F) Box plots for (E) distance from substrate, and (F) *tau*, at the onset of deceleration for feet-contact and head-contact landings. The grey boxes indicate the central 50% data around the median (red line). The whiskers represent 1.5 times the interquartile range. Outliers were included in the analysis. Asterisks represent statistically different comparisons (*, **, ***, and **** represent p<0.05, p<0.01, p<0.001, p<0.0001 respectively). This convention for boxplots and statistical significance is employed for all subsequent figures.

#### Statistical tests

Because we could not *a priori* assume normal distribution of the data on distance from the substrate and *tau* values for the head-contact vs. feet-contact flies, we used a non-parametric (Wilcoxon rank sum) test to compare the various quantities (Fig. 3 E-F; Fig. 4 C-D). All statistical comparisons were performed using MATLAB.

**Fig 4.**
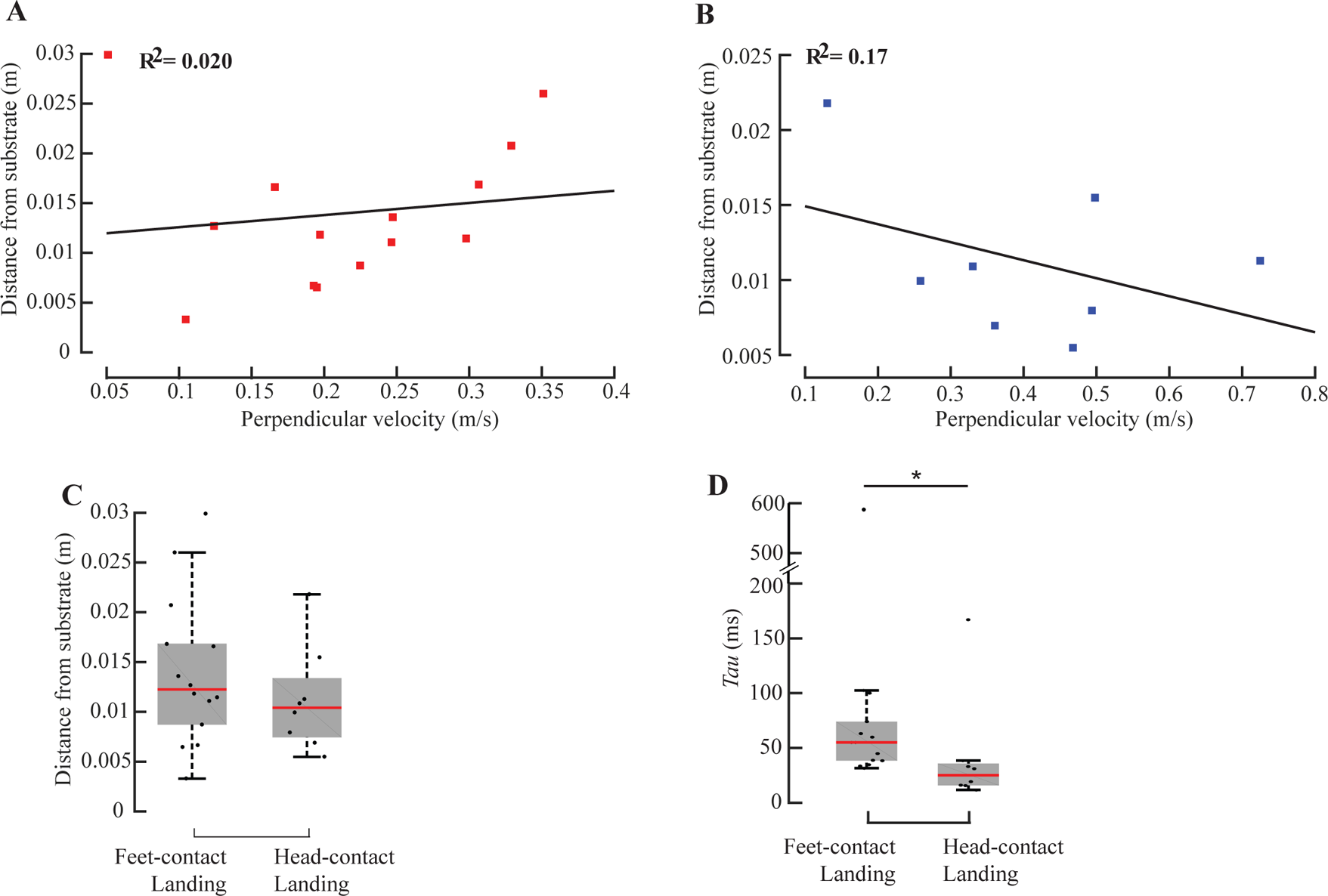
Initiation of leg-extension for flies performing feet-contact and head-contact inverted landings. Of the 22 flies which extended their legs when flying towards the upside down landing platform (see Materials and methods), 14 flies executed a feet-contact and 8 executed a head-contact landing. (A) Distance from substrate versus perpendicular velocity at the time of onset of leg-extension for feet-contact landing trials (n=14; R^2^ = 0.020). (B) Distance from substrate versus perpendicular velocity at the onset of leg-extension for head-contact inverted landing trials (n=8, R^2^ = 0.17). (C-D) Box plots for (C) distance from substrate, and (D) *tau,* at the onset of leg-extension for feet-contact and head-contact landings.

## RESULTS

### Initiation of deceleration and leg-extension before a vertical landing

The landing behaviors for landing on vertical surfaces consist of two components-deceleration of the body and extension of legs-that occur immediately prior to landing. Of the 18 vertical landing trials, we observed a phase of deceleration before touchdown in 13 trials (Fig. 2A). In the remaining 5 trials (Fig. 2B), the flies did not decelerate but we observed leg extension (See Materials and methods). For all cases in which there was a clear deceleration phase, there was a strong linear relationship between displacement and perpendicular velocity at the onset of deceleration (coefficient of determination (R^2^) = 0.71; Fig. 2C). Such flies typically approached the vertical wall at velocities ranging between 0.1-0.55 m/s. These observations support the constant-tau hypothesis for onset of deceleration. In contrast, the correlation between displacement and perpendicular velocity at the onset of leg-extension is weaker (R^2^= 0.17; Fig. 2D), suggesting that leg-extension in landing flies is not initiated at a threshold value of *tau.*

### Initiation of deceleration and leg-extension before an inverted landing

Of the 32 flies which landed on the ceiling, 25 flies decelerated before touchdown (Fig. 2E). In the remaining 7 trials, the flies did not decelerate before touchdown (Fig. 2F) (see Materials and methods). However, they extended their legs. Similar to vertical landings, these flies also typically approached the ceiling at velocities less than 0.4 m/s. For the flies that decelerated, there was only a weak linear relationship between displacement and perpendicular velocity at the onset of deceleration (R^2^= 0.079; Fig. 2G) and at the onset of leg-extension (R^2^= 0.036; Fig. 2H). These results indicate that for inverted landings, neither deceleration nor leg-extension were initiated at threshold values of *tau.*

The inverted landing trials could be grouped into two categories. In 15 trials, the flies bump their head on the landing surface before eventually landing on it, whereas in the remaining 17 trials, the head did not touch the landing surface during the course of the landing maneuver (Fig. 3A). We make the assumption that in the former scenario, which we refer to as *head-contact landing,* flies were unable to land in a controlled fashion, and that the head-on collisions with the landing surface are symptomatic of a lack of control. In the latter scenario, which we call *feet-contact landing,* the flies were able to land with their feet on the surface, and hence we assume that they were in control of their landing maneuver. Flies that landed head-contact into the inverted surface typically approached it with larger perpendicular velocities (blue lines, Fig. 3B) than the flies that landed feet-contact (red lines, Fig 3B).

Out of the 25 flies which decelerated before touchdown (Fig. 2E), 12 performed a feet-contact landing and 13 performed a head-contact landing. The flies that performed inverted feet-contact landings, showed a strong linear relationship between distance from the substrate and perpendicular velocity at the onset of deceleration (n=12; R^2^= 0.69; Fig. 3C), implying that these flies initiated deceleration at a fixed value of *tau.* In flies that landed head-contact, on the other hand, the relationship between the distance from the substrate at which deceleration was initiated vs. perpendicular velocity was weak (R^2^= 0.12; Fig. 3D), suggesting that if a fly does not decelerate at or before the threshold value of *tau,* it is unable to land in a controlled manner. Thus, as shown above, flies initiated deceleration at a constant *tau* before vertical landing, or when landing feet-contact on the inverted surface (Fig. 2C; 3C). However, the correlation between distance from object and perpendicular velocity at the onset of leg-extension was weak, regardless of the type of landing (vertical landing; Fig. 2D; inverted landing; Fig 2H). These results indicate that the deceleration module is elicited independently of the leg-extension module, and perhaps by a different set of cues.

Is there a relationship between the approach kinematics and control of landing? Although the distance from the substrate at which deceleration was initiated was similar for flies that landed feet-contact vs. head-contact (Wilcoxon ranksum test, p>0.05; Fig 3E), there was significant difference in their *tau* values (Wilcoxon ranksum test, p<0.01; Fig 3F). The positive linear relationship for feet-contact landing between distance and perpendicular velocity at the onset of deceleration (constancy of *tau),* and larger values of *tau* at the onset of deceleration of feet-contact as opposed to head-contact suggests that an optimal *tau* margin of 41 ± 9 ms (μ ± σ) was required for initiating deceleration in a properly controlled maneuver. Flies that missed this window were likely to bump their heads against the inverted landing surface. Both the flies that performed feet-contact landings and the ones that landed head-contact, decelerated at rates that do not differ significantly (Wilcoxon Ranksum Test, p>0.05; Fig. S1B), suggesting that flies did not compensate for missing the *tau* margin by increasing average deceleration.

Of the 22 inverted landing trials where the flies initiated leg-extension during flight (and not during take-off, see Materials and methods), 14 executed a feet-contact landing and 8 executed a head-contact inverted landing. The relationship between distance from the substrate and perpendicular velocity at the time of onset of leg-extension was very weak for both feet-contact landing (n=14; R^2^= 0.020; Fig. 4A) and for head-contact landing (n=8; R^2^= 0.17; Fig. 4B). This implies that flies landing feet-contact on inverted surfaces did not initiate leg extension at a constant *tau.* These flies did not significantly differ in the distance from the landing surface at which they began leg-extension (Wilcoxon ranksum test, p>0.05; Fig. 4C), but they began leg-extension at significantly lower values of *tau* as compared to the flies that landed feet-contact (Wilcoxon ranksum test, p<0.05; Fig 4D). This also shows that longer *tau* is essential for landing in a controlled manner.

### Dependence of the initiation of deceleration on the orientation of the landing surface

We next plotted the distance from substrate against perpendicular velocity at the onset of deceleration for both vertical landings (orange) and inverted landings (black; Fig. 5A), and obtained a weak correlation between the two quantities (R^2^= 0.14). These trials however included those flies that landed head-contact. Excluding trials in which the fly landed head-contact, we obtain a stronger correlation between distance from substrate and perpendicular velocity at the onset of deceleration (R^2^= 0.74). This implies that flies that land feet-contact initiate deceleration at the same *tau* before touchdown on both vertical or inverted surfaces (Fig. 5B) and hence the neuronal and mechanistic basis of onset of deceleration may be the same in both cases, regardless of the orientation of the surface.

**Fig 5.**
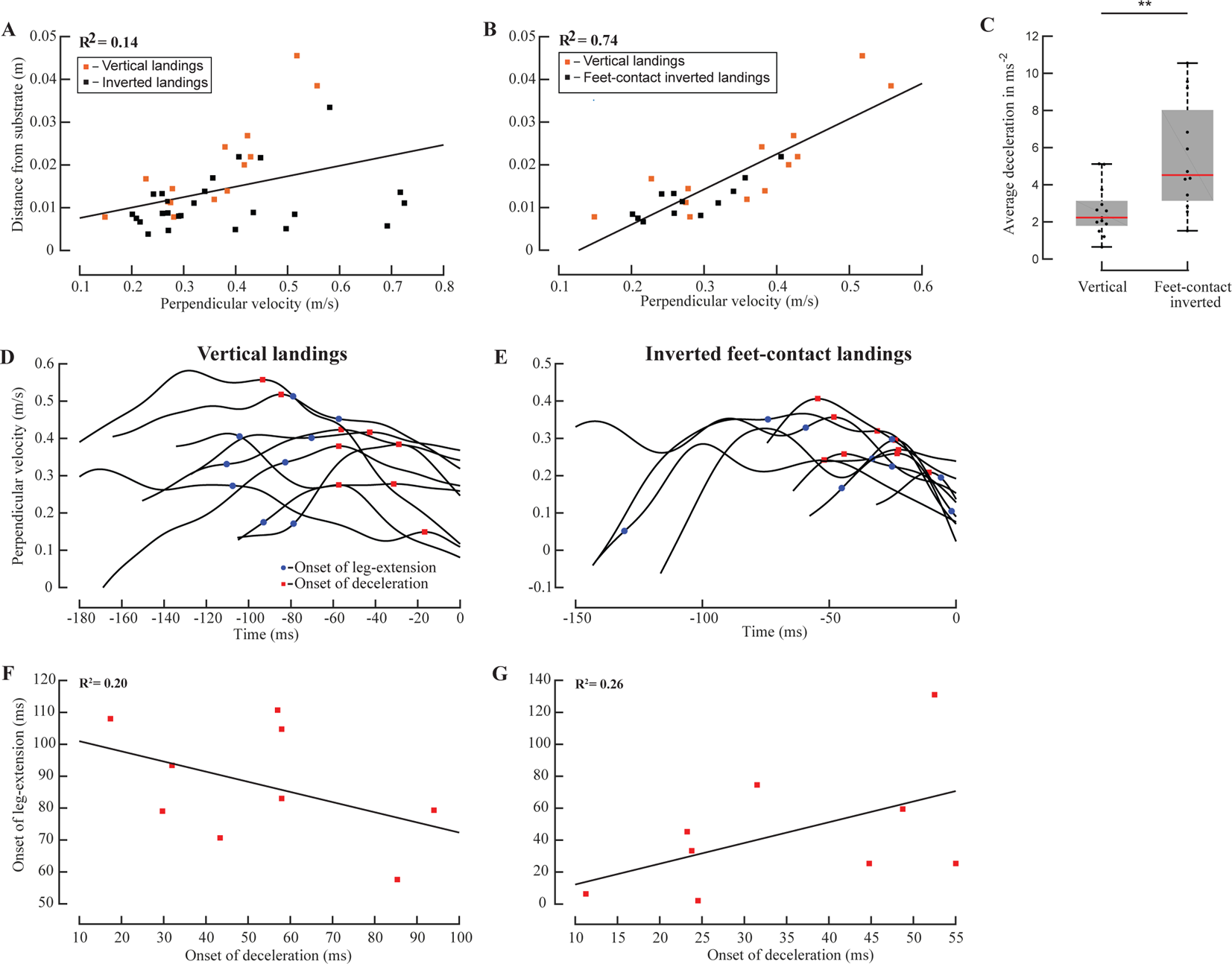
Comparing the onset of deceleration of vertical and inverted landings, and testing for correlation between the onsets of deceleration and leg-extension. (A) Distance from substrate versus perpendicular velocity at the onset of deceleration for vertical (orange squares, n=13) and inverted landings (black squares, n=25, R^2^ = 0.14). (B) Distance from substrate versus perpendicular velocity at the time of onset of deceleration for vertical (n=13) and feet-contact inverted (n=12) landings (R^2^ = 0.74). (C) The flies landing feet-contact on the upside down surface decelerated at significantly higher rates compared to flies landing on the vertical surface (Wilcoxon ranksum test, p<0.01). (D, E) Perpendicular velocity as a function of time for all trials (n=9) in which the onsets of deceleration (red squares) and leg-extension (blue circles) were known (see text for details), for (D) vertical landings and (E) feet-contact inverted landings. (F, G) Time to collision at the onset of leg-extension plotted as a function of time to collision at the onset of deceleration, for (F) all 9 vertical landing trials depicted in (D), and (G) all 9 feet-contact landing trials depicted in (E). The low values of R^2^of best-fit lines show a weak correlation between the quantities on the x and y axes.

Because there is consistency in the onset of deceleration between vertical and inverted feet-contact landings, we wanted to test if a similar stereotypy could be observed in the rate of deceleration. Of all flies that land feet-first on the substrate, those approaching the vertical surface decelerate at lower rates compared to flies approaching the inverted surface (Wilcoxon ranksum test, p<0.01; Fig. 5C). Thus, the rate of deceleration appears to be context dependent.

### Correlation between deceleration and leg-extension

As demonstrated in the previous sections, flies initiated deceleration at a constant *tau* before landing on the vertical surface or smoothly on the upside down surface (Fig. 2C; 3C). However, the correlation between displacement and perpendicular velocity at the onset of leg-extension was weak, regardless of the type of landing (Fig. 2D, H; 4A, B). These results indicate that each module is released by different cues. The subset of trials in which the flies decelerated before touchdown, and in which we could ascertain that leg-extension occurred when flying towards the landing substrate (see Materials and methods), was 9 out of 18 vertical landings, and 9 out of 17 feet-contact landings. Consistency in both the order of initiation of the two modules, and the time difference between the onsets, would support the hypothesis that both modules are initiated by the same set of stimuli. Flies initiated leg-extension *before* deceleration in 7 out of 9 vertical landings (Fig. 5D), and in 4 out of 9 inverted landings (Fig. 5E). Additionally, the correlation between time to collision to the landing surface at the onset of leg-extension and the time to collision to the landing surface at the onset of deceleration was weak for both vertical landings (R^2^= 0.20; Fig. 5F), and feet-contact landings (R^2^= 0.26; Fig. 5G). Therefore, it seems unlikely that the same sensory cues elicit both deceleration and leg-extension.

## DISCUSSION

We filmed houseflies *Musca domestica* landing on a vertical surface (vertical landing) and on the underside of a horizontal surface (inverted landing). Houseflies approaching the vertical surface initiated deceleration at a displacement proportional to the component of flight velocity perpendicular to the landing surface i.e. at a fixed *tau* (Fig. 2C). In nearly half of the flies in the inverted landing assay, there was head-contact while landing (head-contact; see Supplementary videos) whereas the rest touched their tarsi on the surface first before swiveling around and landing (feet-contact) but not their heads. In the case of feet-contact, deceleration was also initiated at a threshold value of *tau* (Fig. 3C). This threshold magnitude of *tau* was similar to the magnitude used by flies while initiating deceleration before touchdown on the vertical surface (Fig. 5B). The correlation between displacement and perpendicular velocity at the onset of leg-extension was weak regardless of the landing surface (vertical or inverted; Fig. 2D, H), or the type of landing (feet-contact or head-contact) (Fig 2H; 4A, B). Flies that performed a head-contact during inverted landings typically approached the landing surface at higher perpendicular velocities (Fig. 3B). Additionally, they triggered both deceleration (Fig 3F) and leg-extension (Fig. 4D) at lower values of *tau* compared to the flies that landed smoothly.

### Computation of *tau* by flies

It has been demonstrated in a previous study that houseflies approaching a sphere initiate deceleration at a threshold value of *tau* (Wagner, 1982). A fly landing on a sphere can potentially contact the surface at any inclination ranging from horizontal to upside down, depending on the orientation of the landing spot which was not recorded in the study. In the current study, we have shown that houseflies initiate deceleration at a fixed value of *tau* regardless of whether they land on a vertical surface, or feet-first on an inverted surface. Visual inspection of our videos of vertical landings reveal that the final moments of the vertical landing maneuver are highly stereotyped: flies always pitch up before contacting with the landing surface (see Supplementary videos). The horizontal velocities of houseflies (Wagner, 1986) and *Drosophila melanogaster* (David, 1978) are known to be inversely correlated with the pitch angle. Therefore, it is likely that flies approaching the vertical surface induce deceleration by increasing their body pitch. However, flies performed inverted landings in a much more variable manner. Such landings involved pitch-up maneuvers prior to landing in some cases, but a combination of roll, pitch and yaw maneuvers before landing in other cases (see Supplementary videos). Despite the variability in the final moments of inverted landings, flies that performed feet-contact inverted landings initiated deceleration at a constant value of *tau.* Moreover, the magnitude of *tau* at the onset of deceleration was also similar for both vertical and inverted landings. Together, these results indicate that flies likely follow the same rules to initiate deceleration before touchdown on any kind of object. A retinal size-dependent threshold model was proposed to explain the initiation of deceleration in *Drosophila melanogaster* approaching a cylindrical surface (van Breugel and Dickinson, 2012). However, the results of the study were experimentally indistinguishable from the constant *tau* model. These results imply that flies can estimate *tau* from optic flow, and initiate deceleration when the value of *tau* falls below a threshold.

The current study adds to the growing body of evidence that nervous systems of animals can compute *tau* and use it to control multiple behaviors. For example, birds approaching a target appear to maintain the rate of change of *tau* with time *(taudot)* at a constant value, resulting in a characteristic deceleration profile (Lee et al., 1991; Lee et al., 1993). Pigeons approaching a perch begin leg-extension at a fixed value of *tau* (Lee et al., 1993). Gannets plunge diving into the sea begin streamlining when the value of *tau* reduces below a threshold (Lee and Reddish, 1981). Bees approaching a surface maintain *tau* at a constant value, resulting in a proportionate decrease in flight velocity with distance (Baird et al., 2013; Srinivasan et al., 2000).

How might flies estimate *tau* from optic flow? When an animal approaches a surface, the instantaneous value of *tau* is approximately equal to the ratio of the angular separation between two points on the surface to the rate of change of angular separation between these two points (provided the points are close in space; Lee, 1976). Thus to estimate *tau,* the nervous system should be able to compute angular size, and rate of angular expansion of objects. Additionally, it must be capable of comparing these two quantities in real time. Despite numerous behavioral examples of *tau* estimation in animals, studies demonstrating neural computation of *tau* are scarce. To the best of our knowledge, the only known example of computation of a threshold value of *tau* by a neuron is in pigeons (Sun and Frost, 1998; Wang and Frost, 1992), which showed that the response onset and peak firing to a looming object of a sub-population of neurons in the nucleus rotundus occurred at a fixed *tau,* irrespective of the angular size or velocity of the object.

Measurement of *tau* can be achieved by comparing the rate of expansion and angular size of a moving stimulus. Are there examples of neurons or neuronal clusters which measure either of these quantities in insects? A recent study in bees revealed descending neurons in the ventral nerve cord monotonically increased their median firing rate with the angular velocity of a frontally presented rotating spiral stimulus, up to a specific angular velocity value beyond which the response saturated. However, the median response of the neurons was also a function of the number of arms in the rotating spiral (which correlates with spatial frequency) (Ibbotson et al., 2017). In flies, the lobula plate tangential cells integrate inputs from local motion detectors and respond to wide-field motion (for a detailed review see Borst et al., 2010). A subset of lobula plate tangential cells called horizontal system (HS) cells respond to optic flow in the horizontal direction (Hausen, 1982). The response of the HS cells to moving gratings depends on the contrast, wavelength, and velocity of the grating (Egelhaaf and Borst, 1989). However, the HS cells of a hoverfly species presented with moving naturalistic images, reliably encoded angular velocity of the images with little dependence on the contrast of the images (Straw et al., 2008). Examples of neurons which measure the angular size of a looming object are seen in animals as diverse as bullfrogs (Nakagawa and Hongjian, 2010), pigeons (Sun and Frost, 1998) and locusts (Gabbiani et al., 1999; Gabbiani et al., 2001). It is thus likely that there may also exist neurons which estimate angular size and angular expansion in the visual neuropil of houseflies. So far, no study has demonstrated neurons which compute the ratio of angular size to angular expansion in insects. A vast majority of the studies of neuronal response to visual stimuli document the firing properties of neurons in the brain or the ventral nerve cord. It is possible that angular expansion and angular size are compared by interneurons in the thoracic ganglia. Studies involving simultaneous presentation of looming stimuli and single unit recordings from the thoracic ganglia are required to test this hypothesis.

A recent study demonstrated that *Drosophila melanogaster* decelerate to a near hover state, followed by acceleration until touchdown on a vertical pole (Shen and Sun, 2017). However, in our study, houseflies decelerated continuously till touchdown in most trials (Fig. 2A, E). Thus, it is likely that there is considerable variation in the visual control of deceleration among flying insects. As mentioned above, houseflies approaching the vertical surface primarily undergo a pitch up maneuver before touchdown. Flies approaching the inverted surface can rotate about all three axes. The biomechanical processes of the landing maneuvers are likely to contribute significantly to the deceleration profile before touchdown, and should be studied in greater detail.

### Variability and versatility of the landing response

15 out of the 32 flies landing on the inverted surface contacted the substrate with their head. Such flies typically approached the ceiling with higher velocity (Fig. 3B), and initiated deceleration and leg-extension at lower values of *tau* (Fig. 3 F; 4 D). This was not observed in flies landing on the vertical surface. Can the differences in experimental setups and procedures for filming vertical and inverted landings (see Materials and methods) explain this observation? For the inverted landing experiments, we illuminated the flight chamber to match the illuminance of sunlight (∼30000 lux). The two halogen lamps used for the purpose did generate considerable heat. Although we turned the halogen lamps on for a maximum of 3 minutes during each trial, we cannot completely rule out the possibility of heat stress affecting the landing behavior.

Crashes into the landing surface have also been documented in previous papers. For instance, around 36% of *Drosophila melanogaster* approaching a cylindrical landing post crashed into it (van Breugel and Dickinson, 2012). In their experiments, the sub-population that crashed did not differ from the landing flies in the retinal size dependent threshold velocity at which they began deceleration. Instead, these flies decelerated at a lower rate, often failing to extend their legs before touchdown. In the current study, we did not find significant differences in the rate of deceleration between feet-contact vs. head-contact landing flies (Wilcoxon ranksum test, p>0.05; Fig. S1B). Also, the head-contact landing flies extended their legs before touchdown, and did not initiate deceleration at a distance proportional to velocity (Fig. 3D). We filmed a single inverted landing from a batch of 4-6 flies. Therefore, we cannot ascertain from our data whether there exists a sub-population of flies are poor at performing inverted landings. It would be interesting to test if the same flies repeatedly bump their heads on the landing surface.

5 out of the 18 flies landing on the vertical surface, and 7 out of 32 flies landing on the inverted surface, did not decelerate before touchdown. It is possible that these flies did not experience sufficiently low values of *tau* to initiate the deceleration response. However, we do not have sufficient number of non-decelerating trials to explicitly test this hypothesis.

For both vertical and inverted landings, flies initiated leg-extension at a point that appears to be independent of distance from the landing substrate, and perpendicular velocity. In 10 out of the 32 inverted landings, the fly initiated leg-extension during takeoff. This implies that either leg-extension is not tightly regulated, or is extremely sensitive to finer cues such as contrast, texture, local light intensities, etc. Indeed, tethered flies initiate leg-extension in response to front-to-back optic flow (Borst, 1986; Borst, 1989; Borst and Bahde, 1986; Borst and Bahde, 1987; Borst and Bahde, 1988b), and the leg-extension response is a function of the size, velocity, and contrast of an object approaching the fly (Borst and Bahde, 1988a; Goodman, 1960). Furthermore, a sudden change in light intensity can lead to leg-extension in tethered flies (Borst, 1986; Goodman, 1960). We are uncertain about the cues that led to the initiation of leg-extension. More studies are required on the leg-extension response in free flight, in which the landing object and the surrounding visual environment are under finer control.

## CONCLUSION

We aimed to understand the rules used by houseflies to initiate two components of the landing maneuver: deceleration, and leg-extension. About half the flies approaching an inverted surface made head-contact before landing. The remaining flies initiated deceleration at a displacement proportional to the component of approach velocity normal to the landing surface. This proportionality constant *(tau)* remained independent of the orientation of the landing surface (vertical or upside down). If a fly missed this *tau* window, it usually contacted its head with the landing surface. The initiation of leg-extension appears to be independent of approach velocity and displacement from the landing surface, indicating that the leg-extension response is either not tightly controlled, or is sensitive to finer cues such as local light intensity changes, body posture, etc.

## ACKNOWLEDGEMENTS

We thank Dinesh Natesan for providing us some of the codes to compute flight variables and Sujeeth Parthiban for his help with the experiments involving vertical landing. We thank all members of the Insect Flight lab for their invaluable inputs on the experimental procedure and data analysis.

## SUPPLEMENTARY FIGURE LEGENDS

**Fig. S1A. Rotation frequency.** We visually estimated the durations of all body rotations in 10 randomly selected videos each of vertical landings and inverted landings. The inverse of the time duration of each rotation is the frequency for the given body rotation.

**Fig. S1B. Comparison of average deceleration between feet-contact and head-contact inverted landings.** Before touchdown, there was no significant difference (Wilcoxon ranksum test, p>0.05) in the rate of deceleration between feet-contact landings and head-contact landings.

## SUPPLEMENTARY VIDEO LEGENDS

**Supplementary Movie 1.** A vertical landing.

**Supplementary Movie 2.** A feet-contact landing.

**Supplementary Movie 3.** A fly bumping onto the ceiling.

**Supplementary Movie 4.** An inverted landing in which the fly pitched up to contact the ceiling.

**Supplementary Movie 5.** An inverted landing in which the fly rolled to contact the ceiling.

**Supplementary Movie 6.** An inverted landing in which the fly rotated about the yaw, pitch and roll axes before touchdown on the ceiling.

